# Progranulin deficiency promotes neuroinflammation through cathepsin B-mediated activation of the NLRP3 inflammasome

**DOI:** 10.64898/2026.10.11.758389

**Authors:** Geetika Aggarwal, Subhashis Banerjee, Ankan Kumar Bhadra, Joel Jihwan Hwang, Don Ashley O. Malabana, Nitya Nagarajan, Denise M. Smith, Jathin Kollarapu, Andrew D. Nguyen

## Abstract

Haploinsufficiency of progranulin (resulting from mutations in the *GRN* gene) is a major genetic cause of frontotemporal dementia (FTD), which is characterized by profound changes in personality and behavior, progressive impairments in executive function, speech, language and/or movement, TDP-43 pathology, lysosomal dysfunction, and neuroinflammation. Studies in progranulin-deficient mouse models have demonstrated the critical role of neuroinflammation in disease pathogenesis, as deletion of NF-kB or complement factors prevents FTD-associated behavioral deficits. While progranulin is known to be anti-inflammatory, the precise molecular mechanism by which its deficiency triggers inflammatory cascades remain unclear. Here, we investigate the involvement of the NLRP3 inflammasome in progranulin-deficient models, including mice, human microglial cells, and patient fibroblasts. In *Grn* mice, we found increased levels of several inflammasome-related proteins, including procaspase-1 and cleaved IL-1β. Similarly, *GRN* knockout (KO) microglia cells showed significant increases in levels of procaspase-1, NLRP3, and cleaved IL-1β. Moreover, KO cells had increased NLRP3 oligomerization, suggesting that progranulin negatively regulates inflammasome assembly. We also investigated the potential link between lysosomal dysfunction and NLRP3 inflammasome activation in progranulin deficiency. We found that pharmacological inhibition of cathepsin B reduced NLRP3 inflammasome oligomerization as well as inflammatory cytokine expression in *GRN* KO cells. Additionally, siRNA knockdown of cathepsin B reduced levels of procaspase-1 and cleaved IL-1β in *GRN* KO cells. Finally, we found that heterozygous *GRN* fibroblasts exhibited similar increases in procaspase-1 and cleaved IL-1β levels, and these increases were corrected by treatment with an antisense oligonucleotide (ASO) that increase progranulin protein levels. Overall, our results demonstrate that progranulin deficiency promotes NLRP3 inflammasome assembly and activity through cathepsin B-mediated signaling, thereby establishing a mechanistic link between lysosomal dysfunction and increased inflammation. The ability of the *GRN*-targeting ASO to correct NLRP3-mediated inflammation highlights a promising therapeutic strategy for FTD-*GRN* and inflammation in other neurodegenerative disorders linked to progranulin deficiency.

## INTRODUCTION

Progranulin is a lysosomal and secreted protein with pleiotropic properties including effects on inflammation, neurite outgrowth, lysosomal function, cell survival, and wound healing [1–6]. Several studies have suggested that full-length progranulin has anti-inflammatory properties, where as its cleavage products, called granulins, are proinflammatory [7–9]. Progranulin has broad neuroprotective effects, with genetic overexpression reducing disease severity in multiple mouse models of neurological diseases, including frontotemporal dementia (FTD) [10–12], Alzheimer’s disease [13, 14], Parkinson’s disease [15], amyotrophic lateral sclerosis (ALS) [16], and ischemic stroke [17]. Similarly, treatment with recombinant progranulin is protective in mouse models of FTD [18], Alzheimer’s disease [19], and traumatic brain injury [20].

Haploinsufficiency of progranulin (encoded by the *GRN* gene) is a major genetic cause of FTD, which is characterized by profound changes in behavior and personality, TDP-43 neuropathology, and neuroinflammation [21–24]. The neuroinflammation includes microglia activation [25–27] which co-localizes with protein aggregation [25] and predicts cognitive decline [26], as well as increased complement C1qa levels detected in the cerebrospinal fluid (CSF) [27]. Additionally, increased levels of astrocyte-derived GFAP have been detected in plasma of individuals with FTD [28, 29], suggesting increased astrogliosis. Cytokine profiling in brain and in blood has suggested possible activation of the NF-kB and/or NLRP3 inflammasome pathways [30]. *Grn* mouse models recapitulate key features of FTD-*GRN* including the behavioral deficits (e.g., decreased sociability, changes in social dominance, and obsessive-compulsive behavior), neuroinflammation, and lysosomal impairment. Notably, progranulin-deficient mice exhibit increased neuroinflammation that includes age-dependent microgliosis and astrogliosis [27, 31–36]. Importantly, studies in *Grn* mouse models have demonstrated the critical role of neuroinflammation in disease pathogenesis, as deletion of NF-kB in microglia [37] or complement factors C1qa globally [27] prevents FTD-associated behavioral deficits.

Studies in microglia and macrophages isolated from *Grn* mouse models have shown that *Grn* knockout cells exhibit a hyper-inflammatory phenotype [33, 36, 38]. For example, stimulation of *Grn* knockout microglia with lipopolysaccharide (LPS) and/or interferon gamma produces markedly higher production of cytokines, including TNF-α, IL-1β, IL-6, and IL-10 [38]. Importantly, these studies demonstrate that progranulin deficiency results in a cell intrinsic, disease-independent effect on inflammation in microglia and macrophages. At the organismal level, *Grn* knockout mice similarly show heightened responses to inflammatory challenges including bacteria-induced pyroptosis [39], LPS-induced endotoxic shock [40, 41], collagen-induced arthritis [9], oxazolone-induced dermatitis [42], and toxin-induced brain injury [38]. Together, these studies indicate that progranulin deficiency primes macrophages, microglia, and possibly other immune cells to mount stronger proinflammatory responses, but the mechanism(s) responsible for these observations are unclear.

The NLRP3 inflammasome is a cytosolic complex that, upon sensing danger signals, activates caspase-1 to drive maturation of the proinflammatory cytokines IL-1β and IL-18 and induce pyroptosis. The NLRP3 inflammasome consists of three proteins: NOD-like receptor protein 3 (NLRP3), apoptosis-associated speck-like protein containing a CARD domain (ASC), and procaspase-1. Assembly of the NLRP3 inflammasome is typically triggered by pathogen-associated molecular patterns (PAMPs) or host-derived damage-associated molecular patterns (DAMPs), which are released in response to injury, cellular stress, or when the immune system is compromised/dysregulated. Once the inflammasome is assembled, procaspase-1 undergoes autocleavage to produce cleaved caspase-1, which subsequently cleaves the proinflammatory cytokines IL-1β and IL-18 from their precursor forms [43–45]. Assembly of the NLRP3 inflammasome and the production of cleaved caspase-1 generally require two signals: priming and activation. The initial priming signal involves the activation of toll-like receptor 4 (TLR4) or cytokine receptors (e.g., TNFR, IL-1R), leading to NF-kB activation and increased expression of inflammatory proteins, including NLRP3, caspase-1, and ASC. Various stimuli can serve as the second signal, including lysosomal dysfunction, mitochondrial dysfunction, endoplasmic reticulum stress, reactive oxygen species production, and exogenous compounds such as crystalline substances (e.g., silica, monosodium urate) and microbial toxins (e.g., nigericin). The leucine-rich repeat (LRR) domain of NLRP3 detects these secondary activation stimuli and leads to ATP-dependent NLRP3 oligomerization, mediated by its NATCH domain. This oligomerization promotes the assembly of NLRP3 oligomers with ASC and procaspase-1 molecules through the N-terminal PYD domain [46–48].

Several recent studies suggest that progranulin deficiency may modulate activity of the NLRP3 inflammasome. In a study with pneumococcal meningitis, *Grn* knockout mice exhibited increased pneumococcal meningitis-induced pyroptosis, as well as increased levels of several NLRP3 inflammasome components in the brain [39]. In another study, progranulin deficiency was reported to decrease NLRP3 inflammasome-mediated inflammation in the context of bone metabolism [49]. It is unclear whether such effects on NLRP3 are linked to progranulin deficiency-mediated lysosomal impairment [35, 50] and/or mitochondrial dysfunction [51, 52], as both are known to serve as signaling cues for NLRP3 inflammasome activation. This study aims to address the mechanisms by which progranulin deficiency modulates NLRP3 inflammasome activation. Our results suggest a model in which progranulin deficiency promotes NLRP3 inflammasome activation, which is in part mediated by activity of the lysosomal protease cathepsin B, which together primes cells to produce hyper-inflammatory responses.

### HIGHLIGHTS

1. The NLRP3 inflammasome is activated in the brain of aged progranulin-deficient mice, as well as in *GRN* knockout human microglia cells
2. This activation of NLRP3 leads to caspase-1-mediated inflammation in progranulin-deficient cells
3. Cathepsin B activity contributes to NLRP3 activation in progranulin-deficient microglia

## MATERIALS AND METHODS

### Mice

*Grn^−/−^* knockout (KO) mice [38] were on the C57BL/6J background (backcrossed more than 8 generations). Mice were housed in a pathogen-free barrier facility with a 12-h light/12-h dark cycle and allowed food and water ad libitum. Pups generated from breeding heterozygous *Grn^+/−^* mice were genotyped by real-time PCR (Transnetyx). All experimental procedures were approved by the Institutional Animal Care and Use Committee at Saint Louis University.

### Cell culture

HMC3 human microglial cells were obtained from American Type Culture Collection (ATCC, CRL-3304) and grown according to the manufacturer’s instructions. HMC3 cells were maintained in Eagle’s Minimum Essential Media (EMEM) (Corning, 10-009-CV) with 10% fetal bovine serum (Gibco, 26-140-095) and 1% penicillin-streptomycin (Gibco, 151-40-122). *GRN^−/−^* KO HMC3 cells were generated by CRISPR/Cas9, as previously described [53]. Human dermal fibroblast (HDF) cells [36] were cultured in Dulbecco’s Modified Eagle Medium (DMEM) (Gibco, 11-995-073) supplemented with 10% fetal bovine serum and 1% penicillin-streptomycin. Cell lines were cultured in 100 mm dishes at 37°C in a humidified 5% CO_2_ incubator.

### Cell treatment, transfection, and transduction

For treatments, HMC3 cells were typically seeded at 5.0×10^5^ cells/well in 12-well plates or at 2.5×10^5^ in 24-well plates. The following day, cells were treated with or without 100 ng/ml LPS (Sigma, L4391) for 4 h. For cathepsin and protease inhibitors, cells (5×10^5^ cells/well in 24-well plates) were incubated with the following inhibitors at 5 μM for 24 h: cathepsin B inhibitor CA-074 methyl ester (MedChemExpress, HY-100350), cathepsin L inhibitor I (Santa Cruz, sc-364671), cathepsin S inhibitor LY3000328 (Sigma, SML3229), and legumain (also called asparagine endopeptidase (AEP)) inhibitor 1 (MedChemExpress, HY-128617). For NLRP3 inhibition, cells (5×10^5^ cells/well in 12-well plates or 1×10^6^ cells/well in 6-well plates) were pretreated with 10 μM of the specific NLRP3 inhibitor MCC950 (InvivoGen, inh-mcc) for 2 h and then primed with 100 ng/ml LPS for 4 h. DMSO and PBS were used as vehicle controls for the respective treatments. For HDFs, cells were seeded at a density of 2.5×10^5^ cells/well in 24-well plates. The following day, cells were treated with or without 100 ng/ml LPS for 3 h, followed by 10 μM nigericin (Sigma, N7143) for a further 15 min.

For siRNA transfection, HMC3 cells were seeded at a density of 5×10^4^ cells/well in 6-well plates. The following day, media was changed prior to transfection with 50 nM siRNA using Lipofectamine 3000 according to the manufacturer’s instructions. After 72 h, cells were treated as described in the figure legends. The following ON-TARGETplus siRNAs were purchased from Dharmacon: non-targeting pool (D-001810-10-50), *GRN* SMARTpool (L-009285-00-0020), and *NLRP3* SMARTpool (L-017367-00-0020).

For lentiviral transduction, HMC3 cells were seeded at a density of 5×10^5^ cells/well in 12-well plates. The following day, the cells in each well were transduced with packaged GFP or mouse progranulin lentivirus at an MOI of 150 in normal culture media. After 24 h, cells were treated with 100 ng/ml LPS or PBS for 4 h.

For ASO treatments, HDF cells were seeded at a density of 2.5×10^5^ cells/well in 24-well plates. The following day, cells were incubated with 5 μM ASO for 24 h. The ASOs used include a non-targeting control ASO and a *GRN*-targeting ASO (M5) that increases progranulin protein levels [54].

### qPCR analysis

Total RNA was isolated from cultured cells using the RNeasy Mini kit (Qiagen, 74106) with on-column DNase digestion (Qiagen, 79256). RNA was reverse-transcribed to obtain cDNA using the iScript cDNA synthesis kit (Bio-Rad, 1708891), and qPCR was performed using PowerUp SYBR Green Master Mix (ThermoFisher, A25777) with a Bio-Rad CFX384 Real-Time System. Primers were synthesized by Integrated DNA Technologies, and the primer sequences are provided in Supplementary Table 1. Results for qPCR were normalized to the housekeeping gene *CYCLO* and evaluated by the comparative C_T_ method.

### Western blot analysis

Cells and flash-frozen tissues were lysed in ice-cold RIPA buffer (50 mM Tris-HCl, pH 7.5, 150 mM NaCl, 0.1% SDS (w/v), 1% NP-40, 0.5% sodium deoxycholate (w/v)) supplemented with protease inhibitors (Roche, cOmplete Mini) and phosphatase inhibitors (Roche, PhosSTOP) and incubated on ice for 10 min. The lysates were centrifuged at 15,000 g for 10 min at 4°C, and the supernatant was saved. Total protein concentration was determined with Bio-Rad BCA Protein Assay. Proteins were electrophoresed on a 4-20% SDS-PAGE gel and transferred onto a nitrocellulose membrane. Membranes were blocked in 5% (w/v) nonfat milk in TBST and incubated overnight with primary antibody at 4°C. After washing, membranes were incubated with HRP-conjugated secondary antibody for 1 h at room temperature. Following washing, the membranes were incubated with SuperSignal West Pico or Femto-enhanced chemiluminescent HRP substrate (ThermoFisher) and visualized using a Chemi-Doc system (Bio-Rad). The following primary antibodies were purchased from abcam: caspase-1 (ab179515), cathepsin D (ab75852), and cathepsin B (ab58802). The following antibodies were purchased from Cell Signalling Technologies: NLRP3 (15101S), cleaved IL-1β (83186S), β-actin (8457S), cathepsin L (71298S), AEP (93627S), and cathepsin S (25084S). Antibodies against human progranulin (AF2420) and mouse progranulin (AF2557) were purchased from R&D Systems. HRP-conjugated secondary antibodies were from Jackson Immuno Research Labs: goat anti-rabbit IgG (111035144), donkey anti-mouse IgG (715035150), and donkey anti-goat IgG (705035147). Data were quantified by densitometry using ImageJ and normalized to β-actin levels.

### NLRP3 oligomerization assay

NLRP3 oligomerization was assessed as described previously [55–57]. Briefly, HMC3 cells were seeded at 5×10^5^ cells/well in 12-well plates. The following day, cells were pretreated with DMSO or specific inhibitors for 1 h, and then treated with 100 ng/ml LPS for 4 h. Cells were lysed in SDD lysis buffer (50 mM Tris-HCl, 150 mM NaCl, 0.5% Triton X-100, 10% glycerol) supplemented with protease inhibitors (Roche, cOmplete Mini) and phosphatase inhibitors (Roche, PhosSTOP). Samples were resuspended in 3X SDD sample buffer (0.5X TBE (Santa Cruz, sc-296650), 2% SDS, 10% glycerol, 0.0025% bromophenol blue) and loaded onto a 1.5% agarose gel. For 1.5% agarose gel preparation, 1.5 g agarose was dissolved in 98 ml of 1X TBE and heated until the gel particles dissolved into a clear solution. 1 ml of 10% SDS was then added, and the gel was cast using a Bio-Rad DNA electrophoretic gel caster. Samples were subjected to agarose gel electrophoresis using 1X TBE containing 0.1% SDS as a SDD running buffer for 30 min at a constant voltage of 130 V. Samples were transferred to a PVDF membrane by wet transfer, and the membrane was used for further western blot analysis as described above.

### Statistical analyses

All statistical analysis was performed using GraphPad Prism 10, and data are presented as means ± SEM. The specific statistical tests used are reported in corresponding the figure legends.

## RESULTS

### Several proteins involved in the NLRP3 inflammasome pathway are increased in the brains of progranulin-deficient mice

Previous studies in aged *Grn^−/−^* mice have found increased neuroinflammation, including age-dependent microgliosis and astrogliosis [27, 31–36]. Therefore, we aged *Grn^−/−^*mice to 18 months and analyzed levels of proteins in the NLRP3 inflammasome pathway in the brain. As expected, heterozygous *Grn^+/–^* and homozygous *Grn^−/−^* KO mice had reduced progranulin protein levels in the cortex and thalamus (Fig. 1). In both tissues, *Grn^−/−^* mice showed a modest increase in caspase-1 protein levels; however, no significant change in NLRP3 levels was observed. In contrast, heterozygous and homozygous *Grn* mice both showed modest increases in IL-1β levels in the thalamus, but no substantial change in IL-1β levels was observed in the cortex (Fig. 1). Together, these results indicate that in aged *Grn^−/−^*mice have elevated levels of several proteins involved in the NLRP3 inflammasome pathway, which is more pronounced in the thalamus.

**Fig. 1.**
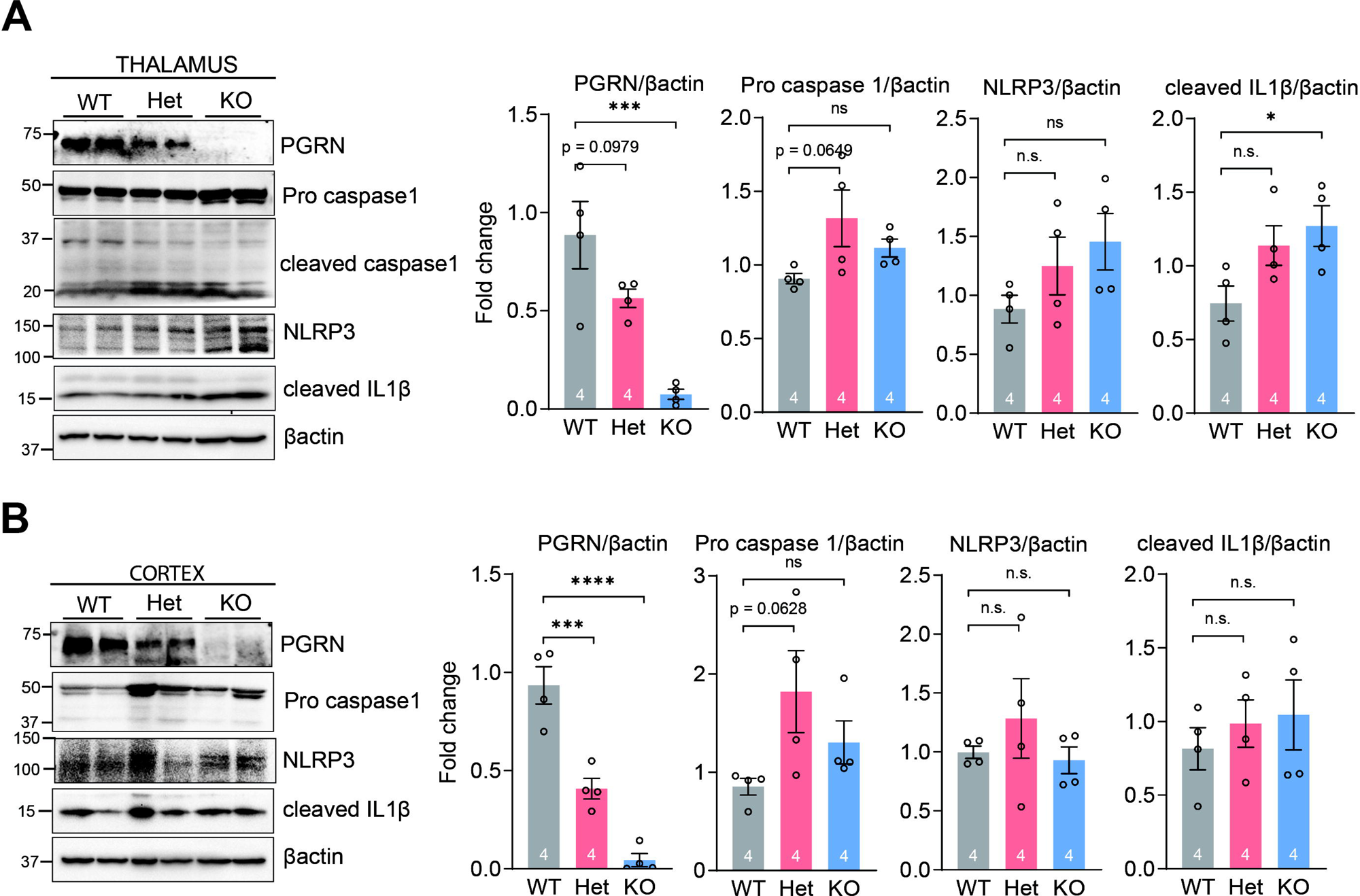
The NLRP3 pathway is activated in brains of *Grn^+/–^* and *Grn^−/−^* mice. Representative western blot analysis of NLRP3 inflammasome-related proteins in the thalamus **(A)** and cortex **(B)** of 18-month-old *Grn* mice (n=4 per group). Graphs show quantification of protein levels normalized to β-actin. Data are presented as mean ± SEM. \**p* < 0.05, \*\*\**p* < 0.001, \*\*\*\**p* < 0.0001, n.s. not significant compared to WT, as determined by one-way ANOVA with Dunnett’s post hoc test. PGRN, progranulin; WT, wild-type; Het, *Grn^+/–^* heterozygous; KO, *Grn^−/−^* knockout.

### Progranulin deficiency increases inflammation in HMC3 human microglia cells

We next used *GRN* KO HMC3 human microglia cells [53] to further investigate the effects of progranulin deficiency on inflammation and NLRP3 inflammasome activation. In unstimulated KO cells, we observed strong increases in the mRNA expression of IL-1α and IL-1β (Fig. 2a), as well as TNF-α and IL-6 [53]. We also observed increased expression of caspase-1 mRNA in KO cells (Fig. 2a). These results in human HMC3 microglia cells are consistent with previous findings of proinflammatory phenotypes in primary mouse microglia cells [38] as well as primary mouse macrophages [33, 36]. To confirm that these changes are attributable to progranulin deficiency, we re-expressed mouse progranulin in the KO cells using lentivirus (Fig. 2b), and that largely corrected the expression of IL-1α, IL-1β, and caspase-1 to levels observed in WT cells (Fig. 2a). These results confirm that the increased expression of these inflammatory genes is due to progranulin deficiency.

**Fig. 2.**
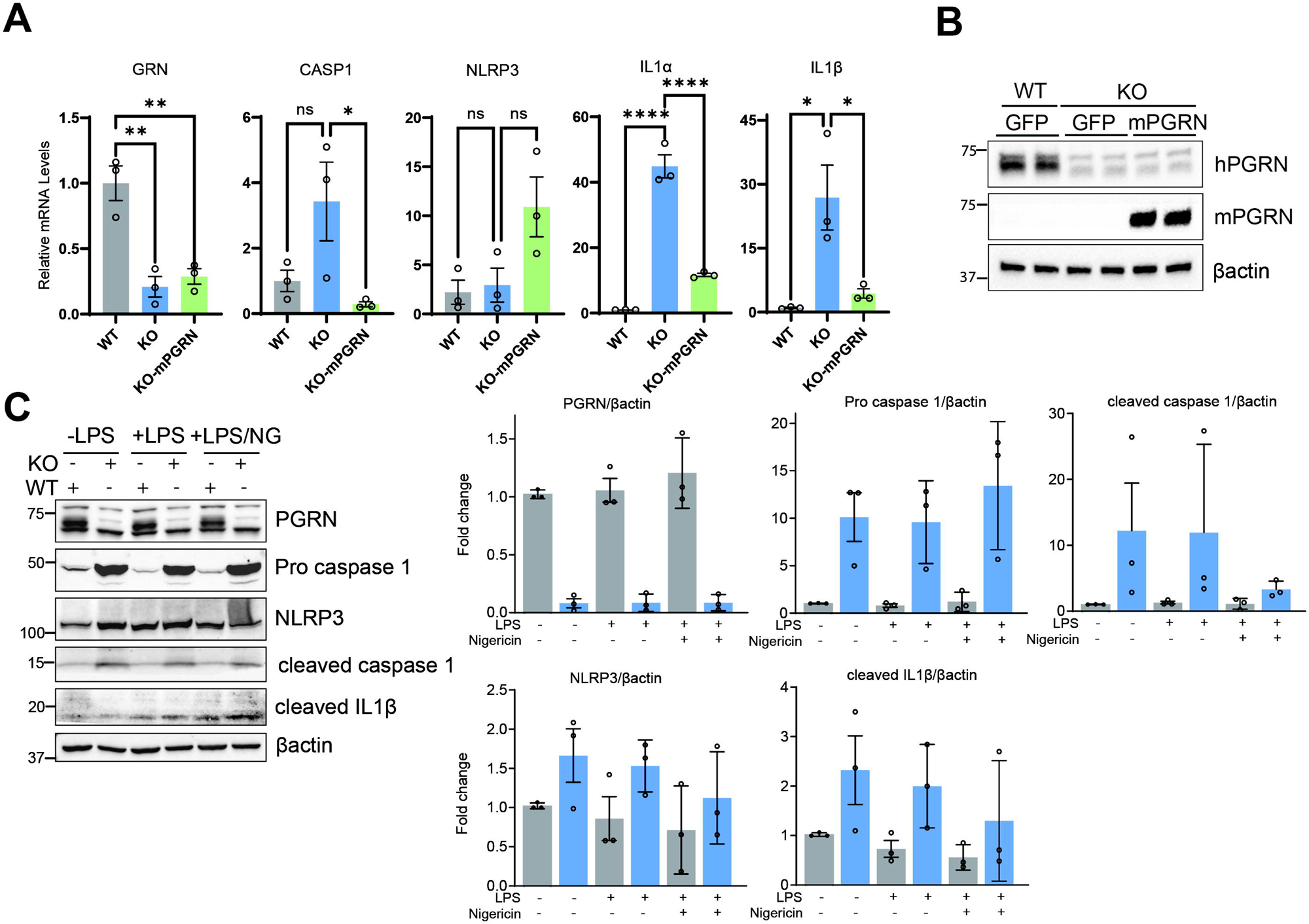
Progranulin deficiency increases levels of NLRP3 inflammasome components and proinflammatory cytokines in HMC3 human microglia cells. **(A)** Relative mRNA expression of GRN, CASP1, NLRP3, IL-1α, and IL-1β in transduced WT vs KO cells. Data are presented as mean ± SEM. \**p* < 0.05, \*\**p* < 0.01, \*\*\*\**p* < 0.0001, n.s. not significant, as determined by one-way ANOVA with Dunnett’s post hoc test. **(B)** Lentiviral expression of GFP control or mouse progranulin (mPGRN) in HMC3 cells, with representative blots confirming expression of mPGRN. **(C)** Representative western blots and quantification of NLRP3, procaspase-1, cleaved caspase-1, and cleaved IL-1β in WT and *GRN* KO HMC3 cells. Cells were treated with either PBS or 100 ng/ml LPS alone or in combination with 10 µM nigericin (NG). Graphs show quantification of protein levels normalized to β-actin. The asterisk (*) indicates a cross-reactive band that is slightly below the progranulin band.

In subsequent studies on NLRP3 inflammasome activation, some groups of cells were treated with the bacterial endotoxin lipopolysaccharide (LPS) to provide the initial priming signal for the NLRP3 inflammasome, and some groups were treated with the bacteria-derived ionophore nigericin (NG) to provide the second activation signal. In the absence of LPS treatment, progranulin deficiency markedly increases procaspase-1, both at the mRNA and protein levels (Fig. 2a, 2c). On the other hand, NLRP3 was modestly increased in KO cells at the protein level, but not at the mRNA level. With LPS treatment, protein levels of procaspase-1 and NLRP3 remained increased in KO cells, and levels of cleaved caspase-1 and IL-1β were also increased (Fig. 2c). These findings reveal that KO cells have increased levels of components of the NLRP3 inflammasome, even under basal conditions, and they suggest that KO cells may have increased activation of the NLRP3 inflammasome.

### Progranulin deficiency causes NLRP3 activation

The NLRP3 inflammasome activation pathway is mediated by two external stimuli/signals. The first signal is activated by DAMPs/PAMPs and accompanied by NF-kB activation [58], which leads to the increased mRNA and protein expression of inflammatory genes including CASP1, NLRP3, and IL-1β. Our findings above in *Grn* mouse brains and *GRN* KO human microglia cells suggest possible increased activation of the NLRP3 inflammasome in the setting of progranulin deficiency. To test this possibility, we knocked down NLRP3 in *GRN* KO HMC3 cells using siRNA and treated cells with LPS. As before, levels of procaspase-1, cleaved caspase-1 and cleaved IL-1β were increased in KO cells (Fig. 3a). Importantly, levels of cleaved caspase-1 and cleaved IL-1β in KO cells were decreased with NLRP3 siRNA, indicating involvement of NLRP3 in IL-1β-mediated inflammation. NLRP3 knockdown also modestly reduced levels of procaspase-1, suggesting that the increased procaspase-1 expression in KO cells is partially controlled by NLRP3 (Fig. 3a).

**Fig. 3.**
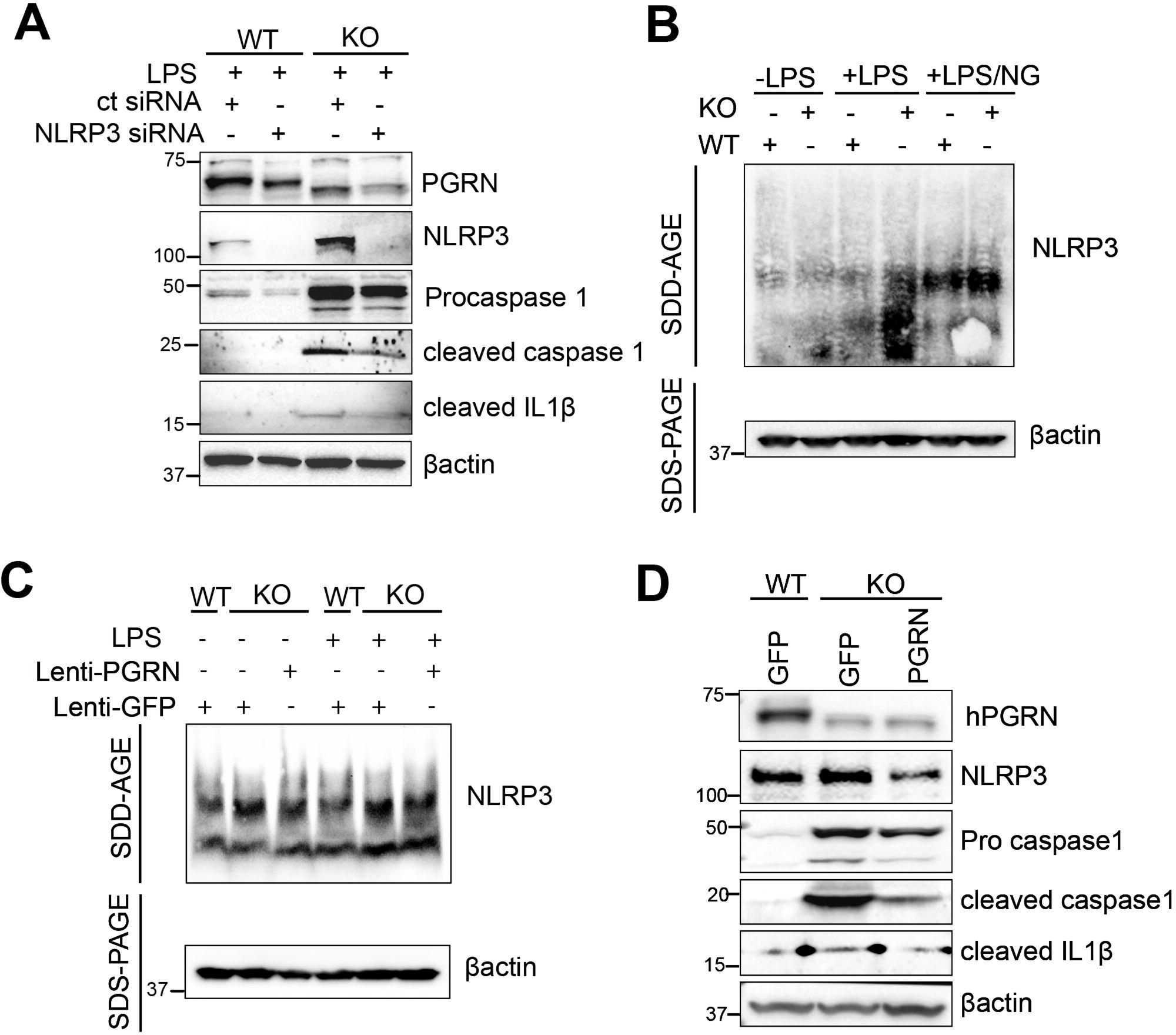
Progranulin deficiency promotes NLRP3 oligomerization and inflammasome activation. **(A)** Representative western blots and quantification of NLRP3, pro- and cleaved caspase-1, and cleaved IL-1β in WT and *GRN* KO HMC3 cells. Cells were transfected with 50 nM of control or NLRP3 siRNA (24 h) followed by LPS treatment (100 ng/ml, 3h); NLRP3 knockdown effectively reduced the elevated levels of inflammasome-related proteins in KO cells. **(B)** SDD-AGE assay of NLRP3 oligomerization in WT and KO cells treated with PBS, LPS (100 ng/ml), or LPS + nigericin (NG, 10 µM). **(C)** SDD-AGE assay showing NLRP3 oligomerization in WT and KO cells (expressing GFP) versus KO cells rescued with mPGRN. mPGRN expression markedly reduced LPS-induced NLRP3 oligomers compared to GFP-expressing KO cells. **(D)** Representative western blots confirming that lentiviral mPGRN expression attenuates the elevated levels of NLRP3, cleaved caspase-1, and cleaved IL-1β in KO cells. The asterisk (*) indicates a cross-reactive band that is slightly below the progranulin band.

Because oligomerization of NLRP3 is a key event in NLRP3 inflammasome activation, we next used semi-denaturing detergent agarose gel electrophoresis (SDD-AGE) to assess NLRP3 oligomerization in WT and KO cells. In the presence of LPS, we found a marked increase in NLRP3 oligomers in KO cells (Fig. 3b), and this effect was reduced by lentiviral re-expression of mouse progranulin in KO cells (Fig. 3c). Lysates from KO cells that received lentiviral expression of mouse progranulin also showed reductions in the levels of NLRP3, procaspase-1, cleaved caspase-1, and cleaved IL-1β, as evaluated by SDS-PAGE (Fig. 3d). Together, these results indicate that progranulin may exert its anti-inflammatory effect through preventing NLRP3 oligomerization and thereby inhibiting activation of the NLRP3 inflammasome.

### Inhibition of cathepsin B partially rescues NLRP3 activation in *GRN* KO cells

To understand the mechanism by which the NLRP3 inflammasome is activated in progranulin-deficient cells, we examined different stimuli that can activate the NLRP3 inflammasome, including lysosomal damage, mitochondrial dysfunction, and changes in ion homeostasis, such as Ca^2+^ and K^+^ [43, 44, 46, 48, 59]. Progranulin deficiency causes lysosomal dysfunction [12, 35, 50, 60], and this is commonly reflected in compensatory upregulation of lysosomal enzymes including cathepsins across many cell types. In agreement with this, we found increased levels of multiple cathepsin enzymes (B, D, L, and S) as well as legumain (which is also called asparaginyl endopeptidase, AEP) in *GRN* KO HMC3 microglia cells (Fig. 4a). Although cathepsin B is known to be involved in NLRP3 activation [61–65], the role of other cathepsins is unclear. Therefore, we tested which cathepsins contribute to NLRP3 activation in progranulin-deficient cells by treating *GRN* KO HMC3 cells with specific inhibitors of different cathepsin enzymes and then measuring effects on IL-1α and IL-1β mRNA levels. We found that only the cathepsin B inhibitor, CA-074 methyl ester, partially reduced (by ∼25-35%) mRNA levels of IL-1α (by ∼35%, p=0.0615) and IL-1β (by ∼25%, p=0.0777) (Fig. 4b). We next tested if the cathepsin B inhibitor treatment reduces NLRP3 oligomerization. For this, we treated KO cells with the NLRP3 inhibitor MCC950 (as a positive control) or the cathepsin B inhibitor CA-074 methyl ester, in the presence of LPS. Both inhibitors reduced levels of NLRP3 oligomers in KO cells, as assessed by SDD-AGE (Fig. 4c). In cell lysates from this experiment, treatment with the cathepsin B inhibitor reduced levels of procaspase-1, cleaved caspase-1, and cleaved IL-1β (Supplementary Fig. 1). To corroborate these findings, we also performed siRNA-mediated knockdown of CTSB. Similar to the results with CtsB inhibitor treatment, CTSB knockdown also decreased levels of procaspase-1, cleaved caspase-1, and cleaved IL-1β protein levels (Fig. 4d), as well as mRNA levels of CASP1, NLRP3, IL-1α and IL-1β (Supplementary Fig. 2). Together, these results show that cathepsin B contributes to the increased activation of the NLRP3 inflammasome and inflammation in *GRN* KO microglia cells, likely initiated by lysosomal impairment.

**Fig. 4.**
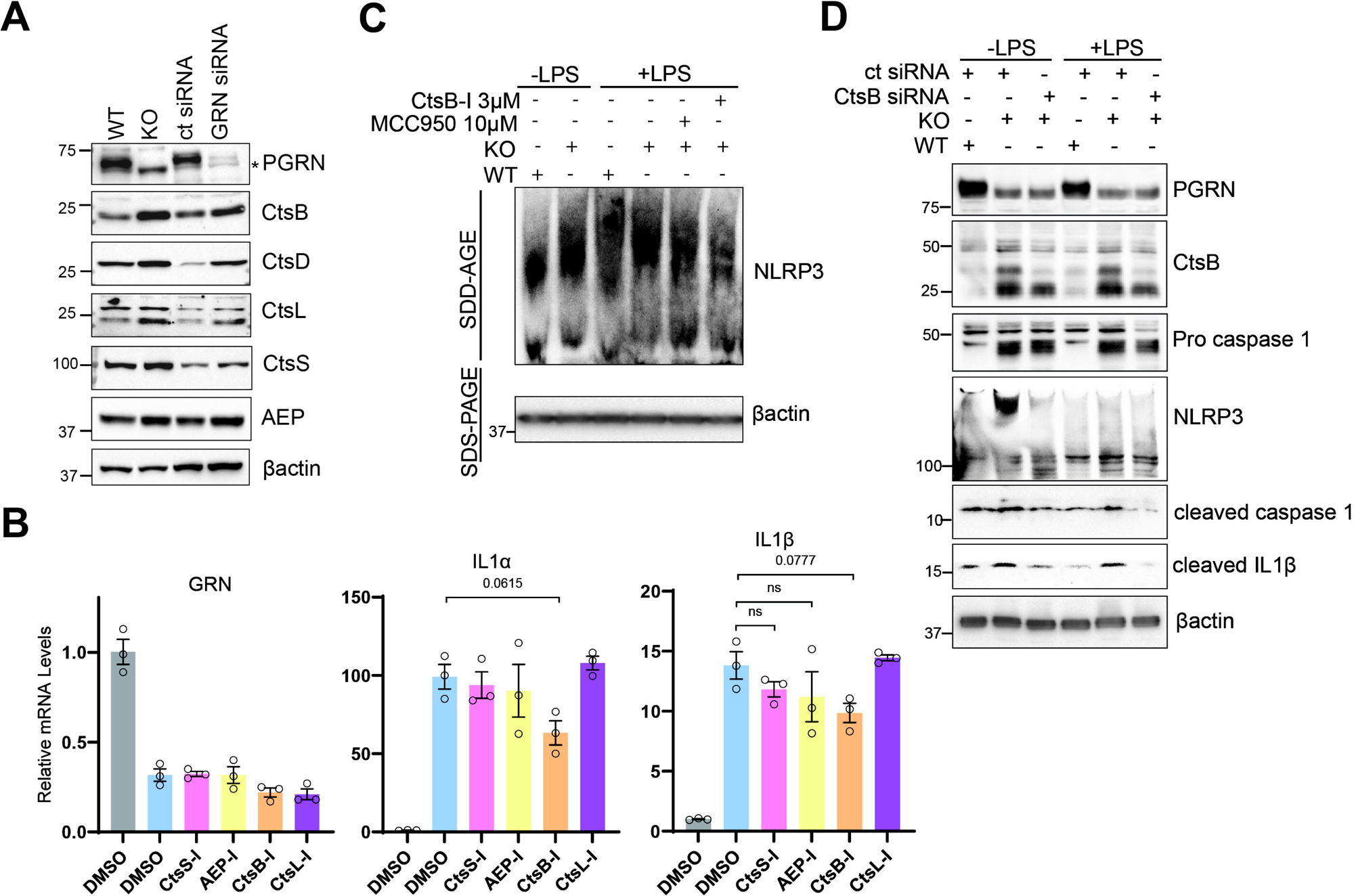
Cathepsin B contributes to increased activation of the NLRP3 inflammasome in *GRN* KO cells. **(A)** Western blots of cathepsin proteins and proteases in HMC3 cells with *GRN* deleted or knocked down by siRNA (50 nM transfection, 24 h). siRNA was performed using 50 nM siRNA for 24 h. **(B)** Relative mRNA expression of GRN, IL-1α, and IL-1β in KO cells treated with DMSO or 5 µM of the indicated cathepsin/protease inhibitors; WT cells treated with DMSO served as the control group. Only the CtsB inhibitor (CtsBi) reduced IL-1α and IL-1β expression. **(C)** SDD-AGE assay to assess NLRP3 oligomerization in WT and KO cells pretreated with DMSO, MCC950 (10 µM), or CtsBi (3 µM) for 2 h, followed by 100 ng/ml LPS treatment for 5h. Both MCC950 and CtsBi reduced NLRP3 oligomer levels in KO cells. **(D)** Representative western blots of NLRP3-related proteins in WT and KO cells. KO cells were transfected with control or CTSB siRNA (50 nM, 24h) prior to LPS treatment (100 ng/ml, 3h). CTSB siRNA effectively attenuated the elevated inflammasome protein levels in KO cells. Data are presented as mean ± SEM. n.s. not significant compared to WT cells treated with DMSO, as determined by one-way ANOVA with Dunnett’s post hoc test. The asterisk (*) indicates a cross-reactive band that is slightly below the progranulin band.

### Progranulin ASO treatment partially rescues NLRP3 activation in *GRN^+/–^*fibroblast cells

We investigated if similar changes in inflammation and the NLRP3 inflammasome are found in *GRN^+/–^* human fibroblasts. For this, we used patient-derived dermal fibroblasts harboring a heterozygous loss-of-function *GRN* mutation (R493X) and control fibroblasts from a family member without the *GRN* mutation [36]. Similar to our findings in *GRN^−/−^*microglia cells, *GRN^+/–^* fibroblasts exhibited increased levels of procaspase-1 and cleaved IL-1β (Fig. 5a).

**Fig. 5.**
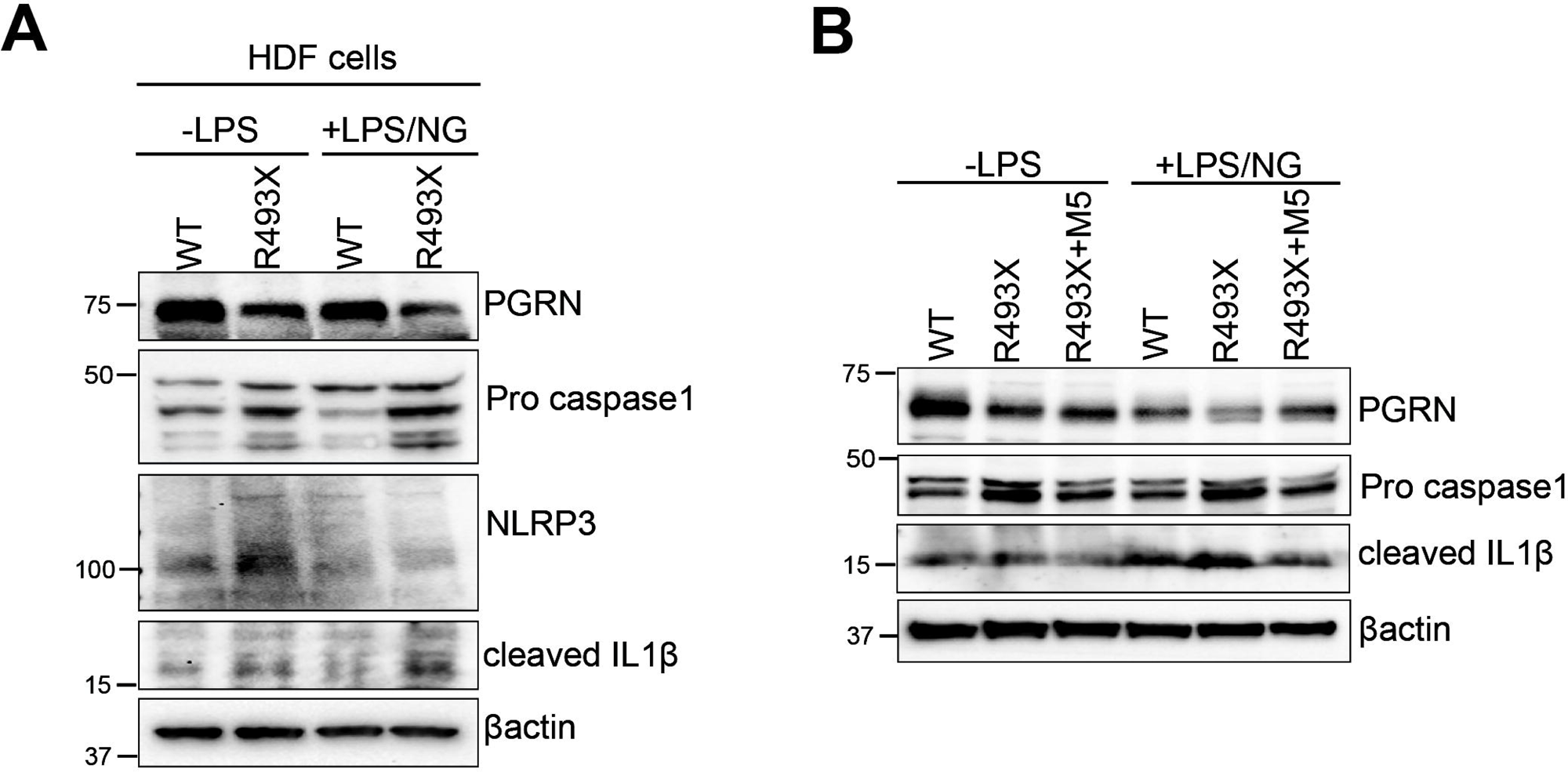
*GRN^+/–^*patient-derived fibroblasts exhibit increased procaspase-1 and cleaved IL-1β levels, which are corrected by pharmacological treatment with an ASO that restores normal progranulin levels. **(A)** Western blot analysis of progranulin, procaspase-1, NLRP3, and cleaved IL-1β in control and *GRN^+/–^*human dermal fibroblasts treated with or without LPS (100 ng/ml) and nigericin (NG, 10 µM). **(B)** Representative western blots of progranulin, NLRP3, and cleaved IL-1β in control and *GRN^+/–^* cells. *GRN^+/–^* cells were treated with a non-targeting control ASO or progranulin-targeting ASO M5 for 48 h prior to LPS stimulation. M5 ASO treatment increased progranulin levels in *GRN^+/–^*cells and attenuated the elevated levels of procaspase-1 and cleaved IL-1β.

To test if pharmacological restoration of progranulin levels can correct the NLRP3 inflammasome hyperactivation in *GRN^+/–^* cells, we treated the fibroblasts with an antisense oligonucleotide (ASO) that increases progranulin levels [54]. This ASO sterically blocks a microRNA binding site in the *GRN* mRNA and thereby increases translation of progranulin protein. As shown in Fig. 5b, ASO treatment increased progranulin protein levels in *GRN^+/–^* cells. Importantly, ASO treatment also corrected procaspase-1 and cleaved IL-1β to levels comparable to those observed in control cells.

## DISCUSSION

We found that proteins involved in NLRP3 inflammasome pathway, including caspase-1 and cleaved IL-1β, are increased in the brain tissue, most pronouncedly in the thalamus, of aged *Grn* heterozygous and homozygous mice. Protein levels of caspase-1, NLRP3, and cleaved IL-1β were also upregulated in progranulin-deficient human microglial cells and in heterozygous patient-derived fibroblasts. Our biochemical studies show that *GRN* KO microglia cells have increased NLRP3 oligomerization, which is a key event in activation of the NLRP3 inflammasome pathway. Using pharmacological inhibitors, we determined that this increased activation of the NLRP3 inflammasome in KO cells is mediated in part by cathepsin B activity, which likely stems from its upregulation due to lysosomal impairment. Importantly, re-expression of progranulin in KO cells reduces NLRP3 inflammasome oligomerization, levels of cleaved caspase-1 and IL-1β, and mRNA levels of proinflammatory cytokine levels.

Because complete progranulin deficiency could have more profound effects including on lysosomal impairment, we also examined effects on the NLRP3 inflammasome in a heterozygous *GRN* system: patient-derived fibroblasts. Similar to our findings in *GRN^−/−^* microglia cells, *GRN^+/–^* fibroblasts exhibited increased levels of procaspase-1 and cleaved IL-1β. This result suggests that haploinsufficiency of progranulin is sufficient to cause increased NLRP3 inflammasome activation. It also indicates that progranulin deficiency activates the NLRP3 inflammasome in multiple cell types, including in non-immune cells. Additionally, in the *GRN^+/–^*fibroblast model, we showed that pharmacological treatment with an ASO that increases progranulin levels can correct the increases in procaspase-1 and cleaved IL-1β levels.

In *Grn* mouse models studies, the thalamus has emerged as a hub of inflammation, including microgliosis and astrogliosis [27, 31–36]. Our results show that caspase-1 and IL-1β are more prominently increased in the thalamus of progranulin-deficient mice as compared to the cortex. In non-pathological settings, caspase-1 is expressed at low levels in its inactive pro-form, procaspase-1. In inflammatory or pathological settings, procaspase-1 levels increase, and when it oligomerizes with another inflammasome protein, NLRP3, it undergoes autocatalysis, generating the cleaved, activated form of caspase-1 that further cleaves pro-IL-1β into the active, proinflammatory IL-1β [43–45, 66]. Our findings of high levels of cleaved caspase-1 in the thalamus of *Grn^+/−^* and *Grn^−/−^* mice suggest upstream activation of the NLRP3 inflammasome. In humans, cytokine profiling in brain and in blood of individuals with FTD has suggested possible activation of the NF-kB and/or NLRP3 inflammasome pathways [30]. Higher levels of caspase-1 and NLRP3 have also been found in other neurological diseases such as Alzheimer’s disease and Parkinsons disease [67, 68]. We note that reduced progranulin levels has been reported to be a risk factor for these diseases [69–77], as well as for limbic-predominant age-related TDP-43 encephalopathy (LATE) [78] and Gaucher disease [79]. While we cannot rule out NLRP3-independent and non-canonical IL-1β pathways in progranulin deficiency, this study primarily focuses on NLRP3 activation in progranulin deficiency.

Microglia are key players in the neuroinflammation aspect of FTD. Studies in individuals with FTD have shown that the neuroinflammation includes microglia activation [25–27], which co-localizes with protein aggregation [25] and predicts cognitive decline [26]. Additionally, studies in *Grn* mouse models have shown that deletion of NF-kB in microglia prevents FTD-associated behavioral deficits [37], highlighting the critical role of microglial inflammation in the disease pathogenesis. To explore whether the upregulated proteins involved in the NLRP3 pathway are due to cell intrinsic effects of progranulin deficiency, we generated a *GRN* KO human microglia cell line. We have observed that, in the absence of LPS, KO microglia are activated with increased expression of proinflammatory cytokines, indicating the effect of progranulin deficiency on the NLRP3 inflammasome pathway. Without LPS, both mRNA and protein levels of caspase-1, NLRP3, and IL-1β are increased. This is in contrast to findings in *Grn* KO mouse microglia and macrophages, which exhibit a hyper-inflammatory phenotype following stimulation with LPS and interferon gamma, but not basal activation [33, 36, 38]. The reason for this difference is not clear, but it could be due to a species-specific effect. Nonetheless, the increased NLRP3 inflammasome activation with progranulin deficiency may help to explain why *Grn^−/−^* mice are more susceptible to a wide range of inflammatory challenges, including bacteria-induced pyroptosis [39], LPS-induced endotoxic shock [40, 41], collagen-induced arthritis [9], oxazolone-induced dermatitis [42], and toxin-induced brain injury [38].

Progranulin regulates lysosomal function, including proteolysis and lipid degradation, and lysosomal impairment is a hallmark feature of progranulin deficiency. This impairment is reflected in elevated lysosomal damage markers such as lipofuscin and galectin-3 [12, 80–82], as well as increased expression of multiple cathepsins in progranulin-deficient cells and tissues [35, 50, 80, 83]. Because cathepsin B interacts with NLRP3 to drive its activation and subsequent inflammation [61–65], it represents a plausible link between lysosomal dysfunction and inflammation. In our experiments testing the role of different cathepsins, only the cathepsin B inhibitor reduced IL-1α and IL-1β mRNA levels; none of the other cathepsin inhibitors tested had an effect. This implicates cathepsin B activity specifically in mediating the inflammation associated with progranulin deficiency. Consistent with this, the cathepsin B inhibitor also reduced NLRP3 oligomerization. Additionally, both pharmacological inhibition and genetic knockdown of cathepsin B reduced levels of procaspase-1, cleaved caspase-1, and cleaved IL-1β in *GRN* KO microglia cells. Together, these findings establish a mechanistic link between lysosomal impairment and increased inflammation in the setting of progranulin deficiency. These perturbations may contribute to the neuroinflammation observed in FTD-*GRN* and potentially also to disease pathogenesis.

## Abbreviations

AEP: asparaginyl endopeptidase
AEPi: asparaginyl endopeptidase inhibitor
ALS: amyotrophic lateral sclerosis
ASC: apoptosis-associated speck-like protein containing a CARD domain
ASO: antisense oligonucleotide
CSF: cerebrospinal fluid
CtsBi: cathepsin B inhibitor
CtsLi: cathepsin L inhibitor
CtsSi: cathepsin S inhibitor
DAMPs: host-derived damage-associated molecular patterns
DMSO: dimethyl sulfoxide
FTD: frontotemporal dementia
HDF: human dermal fibroblast
Het: heterozygous
HRP: horseradish peroxidase
IL: interleukin
KO: knockout
LATE: limbic-predominant age-related TDP-43 encephalopathy
LPS: lipopolysaccharide
LRR: leucine-rich repeat domain
mPGRN: mouse progranulin
NG: nigericin
NLRP3: NOD-like receptor protein 3
PAMPs: pathogen-associated molecular patterns
PBS: phosphate-buffered saline
PVDF: polyvinylidene fluoride
PGRN: progranulin
SDD-AGE: semi-denaturing detergent agarose gel electrophoresis
SDS-PAGE: sodium dodecyl sulfate polyacrylamide gel electrophoresis
TBE: tris-borate-EDTA buffer
TBST: tris-buffered saline with Tween-20
TLR4: toll-like receptor 4
TNF: tumor necrosis factor
WT: wild-type

## Supplementary Information

Supplementary Material 1. Supplementary Table 1. qPCR primer sequences. Supplementary Figure 1. Pharmacological inhibition of cathepsin B attenuates the elevated levels of inflammasome-related proteins in *GRN* KO HMC3 cells. Supplementary Figure 2. Genetic knockdown of cathepsin B partially corrects the increased mRNA levels of inflammasome proteins in *GRN* KO HMC3 cells.

## Acknowledgements

We thank the Genome Engineering & Stem Cell Center at Washington University for generating HMC3 *GRN* KO cell line used in this study and the Hope Center Viral Vectors Core at Washington University for generating lentivirus. We gratefully acknowledge Dr. Heather True (Washington University, St. Louis MO) for providing resources to perform SDD-AGE studies.

## Authors’ contributions

GA and ADN conceived of the study. GA, SB, AKB, and ADN contributed to study design. GA, SB, AKB, JJH, DAOM, NN, DMS, and JK performed experiments. GA, SB, AKB, JJH, DAOM, NN, DMS, and ADN performed data analysis and data interpretation. GA and ADN wrote the manuscript. All authors reviewed and approved the final manuscript.

## Data availability

All data generated or analyzed during this study are included in this published article [and its supplementary information files].

## Declarations

### Ethics approval and consent to participate

Mouse model studies were conducted in accordance with the NIH Guide for the Care and Use of Laboratory Animals and the Institutional Animal Care and Use Committee guidelines for Saint Louis University.

### Consent for publication

Not applicable.

