## Supplementary Information for "Progranulin deficiency promotes neuroinflammation through cathepsin B-mediated activation of the NLRP3 inflammasome"

**Supplementary Table 1.** qPCR primer sequences.

| Human Gene | Forward Primer (5' to 3') | Reverse Primer (5' to 3') |
|---|---|---|
| CASP1 | TTTCTGCTCTTCCACACCAG | TGAAAATCGAACCTTGCGGA |
| CYCLO | GGAGATGGCACAGGAGGAAA | CCGTAGTGCTTCAGTTTGAAGTTCT |
| GRN | ATCTTTACCGTCTCAGGGACTT | CCATCGACCATAACACAGCAC |
| IL-1α | TTGAGTCAGCAAAGAAGTCAAG | GTGGGCCATAGCTTACATGAT |
| IL-1β | GCAGGCCGCGTCAGTTGTTG | TCTGTGGGCAGGGAACCAGC |
| NLRP3 | TTCCTGAGGCTGGCATCT | CTTCGGTCCACACAGAAAGT |

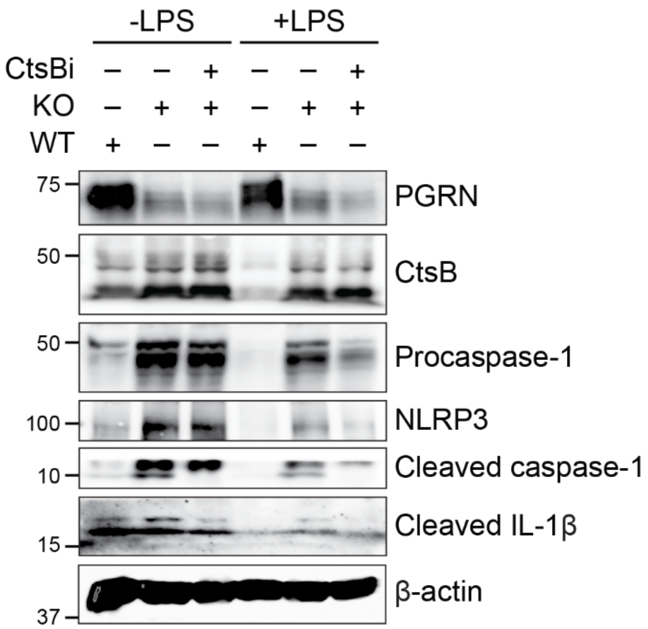


**Supplementary Fig. 1.** Pharmacological inhibition of cathepsin B attenuates the elevated levels of inflammasome-related proteins in *GRN* KO HMC3 cells. Representative western blots of NLRP3-related proteins in *GRN* KO HMC3 cells treated with CtsB inhibitor (CtsBi). KO cells were pre-treated with DMSO or 3 μM CtsBi (2 h) prior to LPS treatment (100 ng/ml, 3 h).

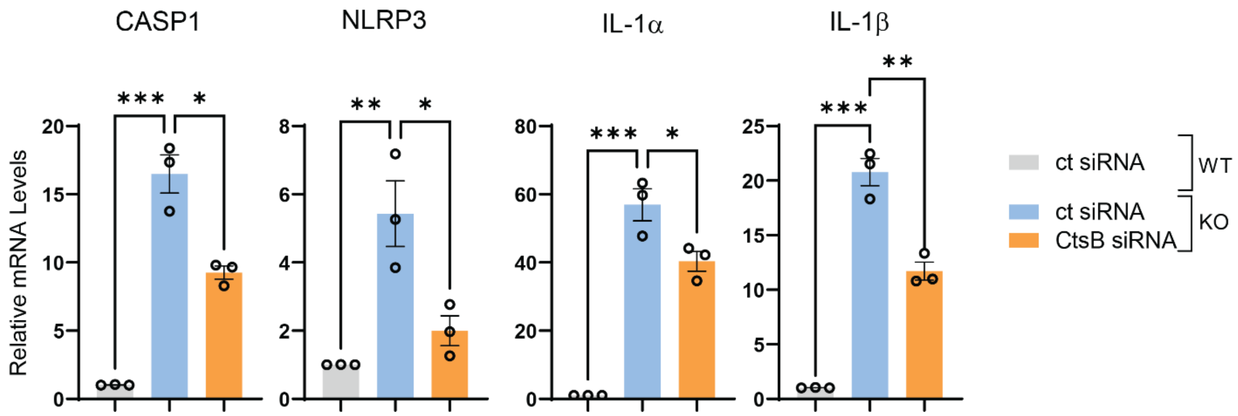


**Supplementary Fig. 2.** Genetic knockdown of cathepsin B partially corrects the increased mRNA levels of inflammasome proteins in *GRN* KO HMC3 cells. Relative mRNA expression of CASP1, NLRP3, IL-1α, and IL-1β in WT vs KO cells following siRNA treatments (control or CtsB). Data are presented as mean ± SEM. *$p < 0.05$, **$p < 0.01$, ***$p < 0.001$, as determined by one-way ANOVA with Dunnett’s post hoc test.
